# Decoupling Efficacy from Toxicity: Engineering Spatial Control in AAV-Mediated Gene Therapy

**DOI:** 10.64898/2025.12.26.696588

**Authors:** Ying Fan, Keqin Tan, Huan Chen, Xiaoqu Chen, Yue Pan, Yajing Chen, Yangsiqi Ao, Ye Bu, Huapeng Li

## Abstract

Hepatotoxicity poses a critical safety challenge for AAV-mediated gene therapy. To mitigate this, we evaluated strategies to minimize off-target hepatic transduction using an antibody expression model. We compared (i) muscle-restricted wild-type AAV9 expression and (ii) a novel myotropic capsid variant, AAV.eM. In humanized B-NDG mice bearing Raji-Luc lymphomas, intravenous administration of AAV9-MHCK7 encoding an αCD19αCD3 bispecific T-cell engager failed to reduce tumor burden. Conversely AAV.eM-MHCK7-αCD19αCD3 substantially alleviated tumor burden and achieved lymphoma clearance. By leveraging tissue-specific microRNAs, precise restriction of AAV.eM-mediated transgene expression to skeletal or cardiac muscle was achieved. Incorporating a heart-specific miR-208a binding site into the transgene’s 3’UTR did not compromise therapeutic efficacy when delivered via AAV.eM-MHCK7. Intramuscular delivery of AAV9-MHCK7-αCD19αCD3 or AAV.eM-MHCK7-αCD19αCD3 both cleared Raji-Luc tumors at a dose of 5 × 10^12^ vg/kg, underscoring the advantage of localized and targeted rAAV delivery over systemic administration. Notably, only AAV.eM-MHCK7-αCD19αCD3 achieved tumor eradication at a tenfold lower intramuscular dose (5 × 10^11^ vg/kg), reducing manufacturing costs and risks of dose-dependent immunogenicity and toxicity. Our findings demonstrate that combining tissue-specific targeting—via engineered capsids or tissue-selective promoters—with local delivery robustly reduces off-target hepatic expression, providing a strategic framework for enhancing the safety of AAV-based gene therapies.

## Introduction

Systemic administration via IV injection of high-dose recombinant adeno-associated virus (rAAV) has been widely adopted as an efficient gene therapy regimen. IV injection of rAAV facilitates widespread transduction of multiple organs, including the central nervous system (CNS) and muscle ^1–3^. A substantial fraction of AAV accumulates in the liver upon entering circulation, particularly for serotypes such as AAV8 and AAV9 ^4, 5^. This liver tropism can be exploited for the treatment of liver-specific diseases such as Crigler-Najjar Syndrome and other metabolic disorders ^4, 6–8^. However, it also increases the risk of hepatotoxicity and, in worst-case scenario, may lead to fatal outcomes in clinical applications. Elevations in alanine aminotransferase (ALT) levels were reported in over 85% of patients who received a single infusion of liver-directed AAV treatment for severe hemophilia A ^9^. Real-world data have documented hepatotoxicity as one of the main adverse events following administration of Onasemnogene abeparvovec (Zolgensma), an AAV9-based gene therapy for spinal muscular atrophy (SMA) ^10^. In 2022, Novartis acknowledged the death of two patients from acute liver failure (ALF) following treatment with Zolgensma ^11^. Four patients with X-linked myotubular myopathy (XLMM) developed cholestatic liver failure and died in the AAV8-based ASPIRO trail ^12, 13^. As of November 2025, three reports of fatal ALF have been documented following treatment of patients with Sarepta Therapeutics’ delandistrogene moxeparvovec-rokl (Elevidys), an AAVrh74-based gene therapy for muscular dystrophy, or an investigational gene therapy using the same serotype ^14^.

The hepatotoxicity associated with rAAV therapy is multifactorial and can be attributed to several mechanism: i) immune-mediated destruction of transduced hepatocytes, notably T cell cytotoxicity in response to the rAAV vectors ^15, 16^; ii) direct liver injury caused by viral capsid components or transgene over expression ^17^; iii) vector overload-induced cellular stress, endoplasmic reticulum stress, and apoptosis ^9, 18^; iv) pre-existing liver conditions ^19^. Growing concerns have been raised in the scientific community regarding potential hepatotoxicity associated with rAAV-based gene therapies underscore the need for comprehensive safety evaluations. Accordingly, the development of strategies to achieve tissue-restricted transgene expression using wild-type AAV or the engineering of tissue-specific AAV capsid variants with minimized liver tropism, is of paramount importance for mitigating hepatotoxicity in rAAV-mediated therapeutic interventions.

In this study, we conducted a comparative evaluation of multiple AAV-based delivery strategies to circumvent off-target expression, using *in vivo* production of an αCD19αCD3 bispecific T cell engager (BiTE) as a model system. We benchmarked our novel myotropic capsid variant, AAV.eM—engineered by inserting an RGD-containing peptide into an innovative backbone with globally reduced tissue tropism and minimized liver targeting ^20^—against several approaches designed to reduce the high-level hepatic transgene expression associated with AAV9.

## Results

### Systemic administration of AAV.eM outperforms AAV9 combined with muscle-specific promotor

To circumvent hepatotoxicity in rAAV-mediated gene therapy, we implemented two distinct approaches: (i) deployment of the novel engineered AAV.eM capsid variant, which exhibits liver-detargeting and muscle-tropic properties, for transgene delivery under the control of the muscle-specific MHCK7 promoter; and (ii) utilization of the wild-type AAV9 serotype coupled with the same MHCK7 promoter to achieve muscle-restricted transgene expression. We evaluated the therapeutic efficacy of both strategies in humanized immunodeficient B-NDG mice bearing Raji-Luc lymphomas (Fig. 1A). As a benchmark control, we included a treatment group receiving AAV9-delivered transgene driven by the ubiquitous CAG promoter. We selected the well-characterized bispecific T cell engager αCD19αCD3 as the transgene and compared the therapeutic efficacy of AAV.eM to that of AAV9 in humanized immunodeficient B-NDG mice bearing Raji-Luc tumors.

**Fig. 1.**
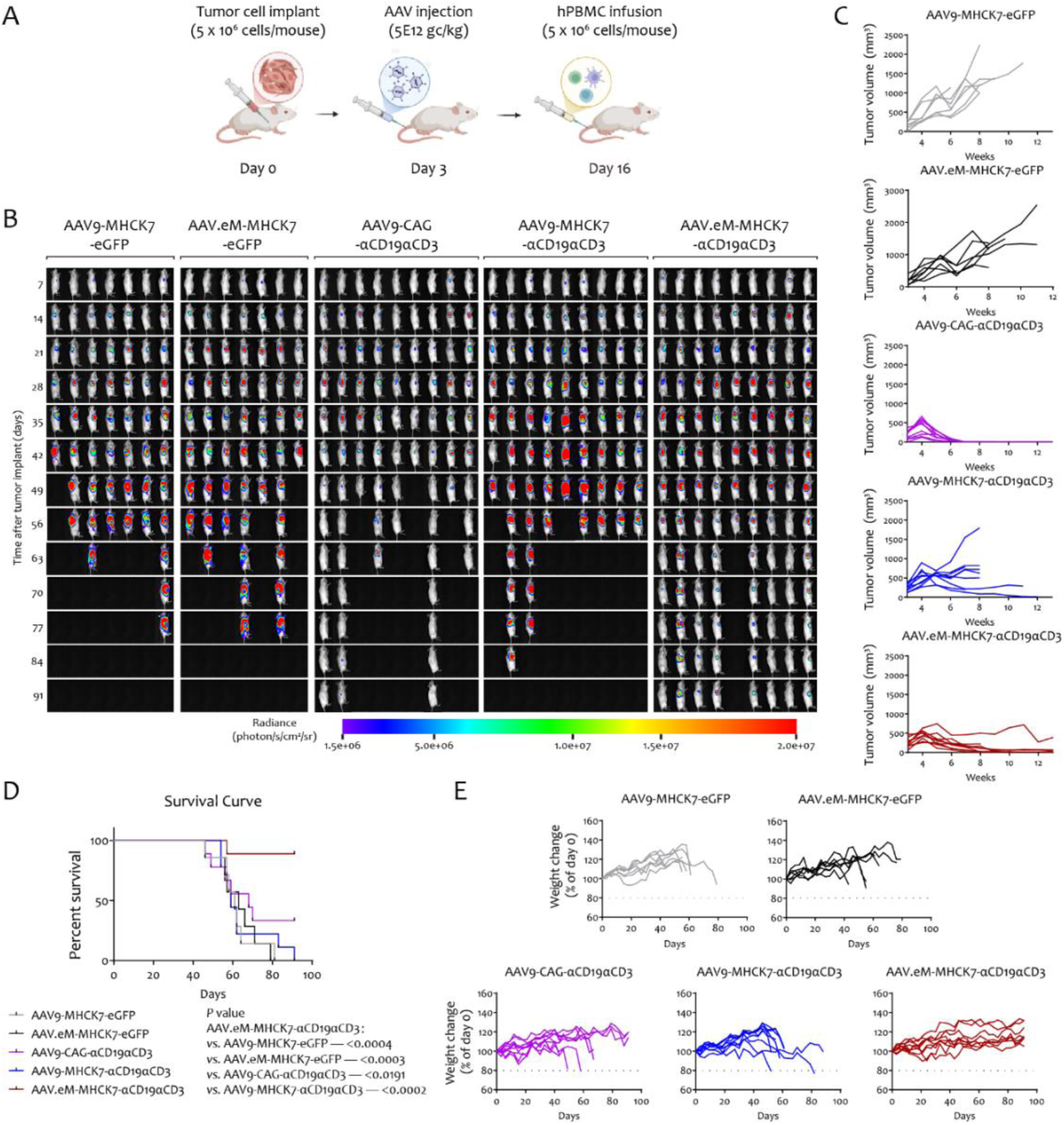
Evaluation of intravenously administered, AAV-vectored bispecific antibody therapy in a humanized lymphoma model. **(A)** Schematic of the experimental timeline and treatment regimen. **(B)** *In vivo* bioluminescent imaging of Raji-Luc-tumor-bearing mice in different treatment groups over time. AAV9-MHCK7-eGFP and AAV.eM-MHCK7-eGFP are control groups expressing eGFP. AAV9-CAG-αCD19αCD3, AAV9-MHCK7-αCD19αCD3, and AAV.eM-MHCK7-αCD19αCD3 are therapeutic groups expressing αCD19αCD3, a bispecific T-cell engager composed of single-chain antibodies for CD19 and CD3 joined by a peptide linker. CAG denotes a cyto-megalovirus (CMV) early enhancer/chicken β-actin promoter, MHCK7 denotes a muscle-specific promoter, and AAV.eM refers to an engineered AAV capsid with reduced liver tropism and enhanced muscle tropism. [n = 10 for AAV9-MHCK7-eGFP treated mice. n = 10 for AAV.eM-MHCK7-eGFP treated mice. n=13 for AAV9-CAG-αCD19αCD3 treated mice. n=13 for AAV9-MHCK7-αCD19αCD3 treated mice. n=13 for AAV.eM-MHCK7-αCD19αCD3 treated mice. n= 3-4 from each group were used for the detection of αCD19αCD3 BiTE, cytokines and chemokines in the serum.]. **(C)** Tumor growth kinetics measured by caliper. Tumor volume was tracked over time in mice treated with the indicated AAV. **(D)** Kaplan-Meier survival curves for each treatment cohort. Statistical significance between groups was assessed with the Log-rank test. P value was shown in the panel. **(E)** Individual mouse body weights presented as percent change from baseline, monitored throughout the study period.

To characterize *in vitro* production and secretion of functional BiTE molecules, an AAV9 viral vector encoding the amino acid sequence of the clinically relevant blinatumomab was packaged and used to transduce HEK293T cells (Supplementary Fig. 1A). The secreted BiTEs exhibited specific and concentration-dependent binding to their cognate antigens. This was demonstrated by a competitive binding assay, in which cell culture supernatant effectively competed with fluorescently labeled anti-CD3ε and anti-CD19 antibodies for binding to Jurkat and Raji cells, respectively (Supplementary Fig. 1B). Similar findings were obtained with a commercially available blinatumomab equivalent. Specificity was further confirmed by the lack of competition from supernatant of HEK293T cells transduced with a control AAV9-eGFP vector. Functional efficacy of the AAV-expressed BiTEs was demonstrated through the induction of classic T-cell effector functions. Co-culture experiments resulted in potent, T-cell-dependent cellular cytotoxicity, as quantified by the significant release of lactate dehydrogenase (LDH) and an increase in Annexin V staining in target cell populations (Supplementary Fig. 1C). Furthermore, engagement of the TCR complex by the BiTE molecules triggered robust activation of naïve CD3+ T cells, evidenced by the substantial secretion of IL-2 and IFN-γ (Supplementary Fig. 1D). Collectively, these data establish that AAV-mediated gene delivery of the αCD19αCD3 BiTE construct enables the efficient production and secretion of fully functional BiTE molecules capable of directing T-cell-mediated cytotoxicity and potently activating T-cell immune responses *in vitro*.

A single tail vein injection of AAV.eM-MHCK7 expressing αCD19αCD3 BiTE resulted in superior tumor growth control and prolonged survival compared with the AAV9-MHCK7 approach, as assessed by *in vivo* imaging, caliper measurements, and survival monitoring (Fig. 1, A-D). The AAV.eM-MHCK7 treatment group exhibited significant tumor regression, although complete eradication was not achieved, possibly due to delayed humanization (hPBMC infusion initiated on day 16 post Raji-Luc tumor implantation in this experiment). In contrast, tumor progression in the AAV9-MHCK7 group was essentially uncontrolled relative to eGFP-treated controls, resulting in high mortality. While AAV9-CAG-mediated delivery of αCD19αCD3 BiTE suppressed lymphoma growth, leading to progressive tumor reduction and eventual clearance, this group exhibited unexpected mortality despite evident therapeutic benefit (Fig. 1B). Mortality coincided with rapid body weight loss in AAV9-CAG-αCD19αCD3-treated mice prior to death (Fig. 1E).

A subset of mice underwent terminal blood collection on day 28 for quantification of serum αCD19αCD3 BiTE. We observed a strong correlation between serum αCD19αCD3 BiTE levels and effective tumor growth inhibition across treatment groups (Fig. 2A). Serum concentrations of αCD19αCD3 BiTE in the AAV.eM-MHCK7 group were approximately 2.5-fold higher than those observed in the AAV9-MHCK7 group. Notably, the AAV9-CAG-αCD19αCD3 group exhibited the highest serum drug concentration, which was associated with its superior tumor growth suppression and clearance efficacy. To assess treatment-related toxicity, day 28 serum was also evaluated for hepatic and inflammatory markers (Fig. 2B-C). No significant differences were observed for most parameters, except for CCL2 and CCL5, which were significantly higher in the AAV9-MHCK7-eGFP control group compared with other control and treatment groups.

**Fig. 2.**
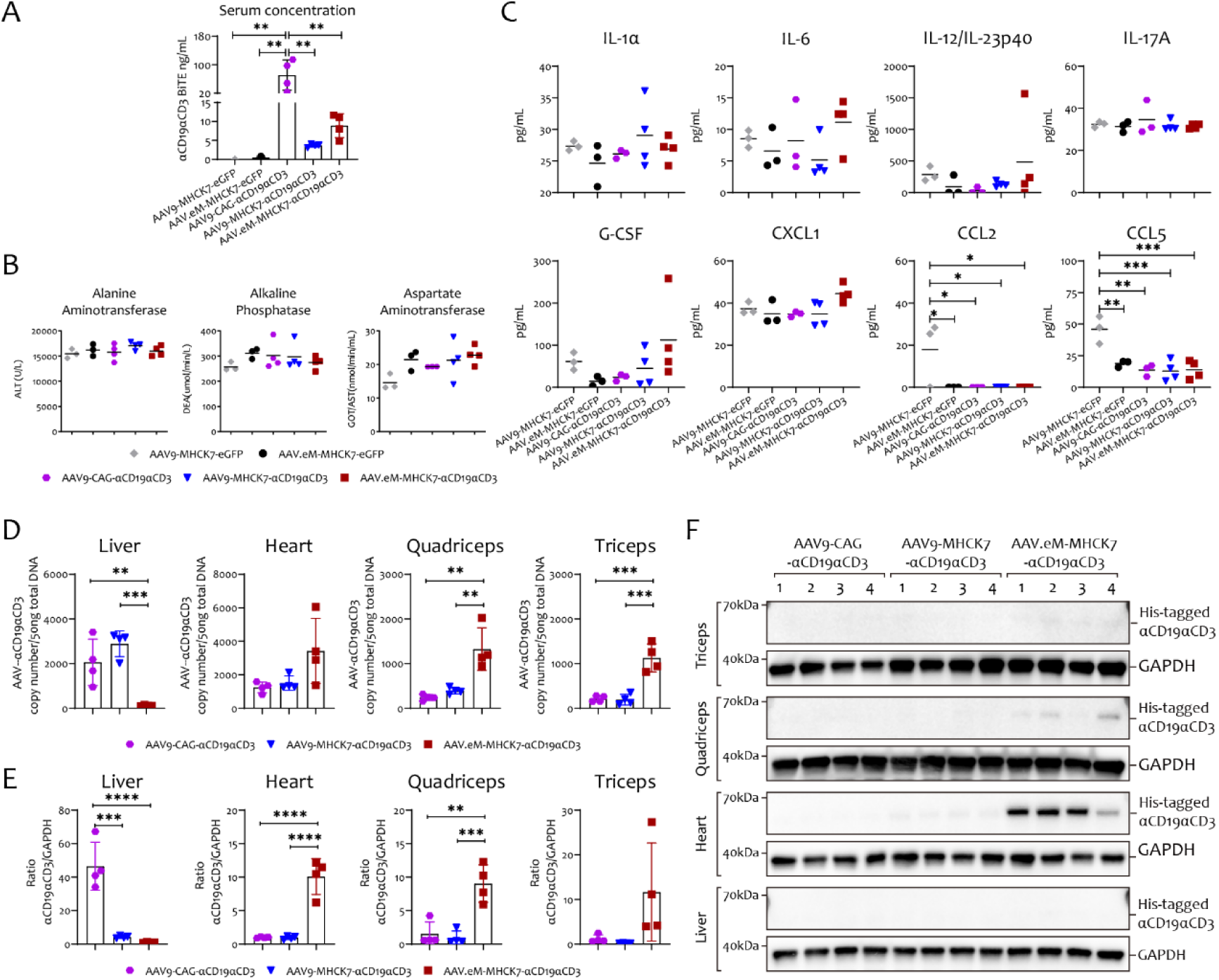
Pharmacokinetic, safety, and biodistribution profile of intravenously administered AAV-encoded bispecific antibody in a humanized lymphoma model. Mice were treated per the regimen in Fig. 1 and euthanized on day 28 post-engraftment for analysis. **(A)** Serum concentration of the AAV-expressed anti-CD19CD3 BiTE was quantified by anti-His tag ELISA. **(B)** Plasma levels of alanine aminotransferase (ALT), alkaline phosphatase (ALP), and aspartate aminotransferase (AST). **(C)** Serum concentrations of murine cytokine and chemokine profiles including IL-1α, IL-6, IL-12/IL-23p40, IL-17A, G-CSF, CXCL1, CCL2, and CCL5, were assessed by cytometric bead array (CBA). Values represent the mean ± SD (n=3-4). Statistical significance was determined by one-way ANOVA. * *P*<0.05, ** *P*<0.01, *** *P*<0.001. **(D)** Biodistribution of αCD19αCD3 genome across tissues in AAV9-CAG-αCD19αCD3, AAV9-MHCK7-αCD19αCD3, and AAV.eM-MHCK7-αCD19αCD3 therapeutic groups. Tissue samples collected on day 28 post tumor cell implantation. Bar graphs show AAV vector genome copy numbers per 50 ng total DNA in heart, liver, triceps, and quadriceps at study endpoint. Data are presented as mean ± SD (n=3-4). Statistical significance was determined by one-way ANOVA. ** *P*<0.01, *** *P*<0.001. **(E)** Transcript levels of the αCD19αCD3 BiTE in indicated tissues collected on day 28 post tumor cell implantation, determined by quantitative RT-PCR. Bar graphs show quantification of αCD19αCD3 normalized to GAPDH. Data are presented as mean ± SD (n=3-4). Statistical significance was determined by one-way ANOVA. ** *P*<0.01, *** *P*<0.001, **** *P*<0.0001. **(F)** Western blot analysis of His-tagged αCD19αCD3 protein expression in the indicated tissues collected on day 28 post tumor cell implantation. GAPDH was used as a loading control.

Quantitative real-time PCR analysis of AAV genomes revealed pronounced liver tropism for WT AAV9, with both AAV9-CAG and AAV9-MHCK7 exhibiting significantly higher viral genome accumulation compared with AAV.eM-MHCK7 in hepatic tissue (Fig. 2D). As expected, AAV.eM showed substantial accumulation in both cardiac and skeletal muscle tissues (Fig. 2D). Employing the muscle-specific promoter MHCK7 in combination with AAV9 effectively restricted αCD19αCD3 BiTE expression to muscular tissues, as only minimal transcript levels were detected in the liver (Fig. 2E). Consistent with viral genome quantification, AAV.eM-MHCK7 yielded the highest αCD19αCD3 BiTE transcript levels in both cardiac and skeletal muscles, approximately tenfold greater than those mediated by AAV9-CAG and AAV-MHCK7 (Fig. 2E). Although protein signals were relatively weak, higher transgene protein levels were observed in the triceps and quadriceps of the AAV.eM-MHCK7-αCD19αCD3 group compared with other groups (Fig. 2F). Interestingly, higher transgene protein level was detected in the cardiac tissue relative to quadriceps and triceps by western blotting following AAV.eM-MHCK7-mediated delivery of anti-CD19CD3 BiTE (Fig. 2F).

To elucidate the mechanistic basis for the differential expression profiles of αCD19αCD3 BiTE, with significantly higher levels detected in cardiac tissue compared with skeletal muscle following AAV.eM-MHCK7m administration, we intravenously administered AAV vectors carrying dual reporter genes (firefly luciferase 2 and eGFP) to BALB/c mice. Live imaging revealed robust hepatic transduction in the AAV9-CAG group as early as 3 days post-injection, peaking at day 7, indicating predominant viral accumulation and transgene expression in the liver (Supplementary Fig. 2A). In contrast, AAV.eM-MHCK7 exhibited progressive, pan-muscular biodistribution with sustained signal amplification over 6 weeks (Supplementary Fig. 2A). Notably, the AAV.eM-MHCK7 group elicited significantly higher luciferase activity/eGFP signals than AAV9-MHCK7, underscoring the superior muscle transduction efficacy of the engineered AAV.eM capsid (Supplementary Fig.2 A-C). Western blot analysis confirmed robust eGFP protein expression in triceps, quadriceps, and cardiac tissues in AAV.eM-MHCK7-eGFP injected group (Supplementary Fig.2 B), indicating that AAV.eM mediates efficient transgene expression in both cardiac and skeletal muscle. Consistent with the protein data, eGFP mRNA levels were substantially higher in the quadriceps, triceps, and heart of mice administered with AAV.eM-MHCK7-eGFP compared with AAV9-MHCK7, corroborating the enhanced muscle-tropism of AAV.eM (Supplementary Fig. 2C). Together, these findings suggest that the pronounced disparity in αCD19αCD3 BiTE protein levels between cardiac and skeletal muscle tissues in the AAV.eM-MHCK7 group (Fig. 2F) may reflect more efficient secretion of the BiTE from skeletal muscle into the circulation, resulting in lower steady-state tissue retention despite high transcript levels. Similarly, the absence of detectable αCD19αCD3 BiTE in hepatic tissue from the AAV9-CAG group is likely due to secretion of the protein into the extracellular space (Fig. 2F).

### MicroRNA-mediated tissue-restricted transgene expression

AAV.eM exhibits general muscle tropism without fine specificity for distinct muscular compartments. For therapeutic applications requiring spatial precision, a strategy beyond the canonical use of tissue-specific promoters such as MCK or cTnT is desirable. Here, we posited that 3’ UTR engineering with tissue-selective microRNA response elements could provide an additional layer of control. By harnessing distinct endogenous microRNA patterns, this approach can spatially restrict transgene expression to either myocardial or skeletal muscle compartments, thereby enabling highly precise, disease-specific targeting.

We employed miR-208a and miR-206 as tissue-specific microRNAs capable of discriminating cardiac muscle from skeletal muscle ^21, 22^. The liver-specific microRNA miR-122 was incorporated as a positive control of microRNA mediated knockdown ^23^. Three tandem repeats of microRNA binding sites were cloned into the 3’ UTR of the target gene for miR-122 and miR-206. For robust recruitment of miR-208a, which is expressed at relatively low absolute levels in the heart ^24^, five binding site repeats were inserted into the 3’UTR (Fig. 3A). To assess the efficacy and specificity of microRNA-mediated tissue-specific knockdown, BALB/c mice were systematically injected with AAV carrying an eGFP report gene harboring microRNA recognition sites downstream of the coding sequence. Imaging of whole tissues isolate at day 9 and day 21 post-infection demonstrated that this strategy successfully restricted target gene expression in a tissue specific manner, selectively silencing expression either in the heart or in skeletal muscle, depending on the microRNA used (Fig.3B). Furthermore, to determine the minimal number of miR-208a binding sites required for efficient cardiac-specific knockdown without causing excessive sequestration of miR-208a, we generated constructs harboring one to five binding site repeats in the 3’UTR of eGFP. This was also prompted by an observed reduction of eGFP signal in skeletal muscle at day 9 post-infection in mice injected with AAV containing five miR-208a binding sites, suggesting very low-level miR-208a expression in the skeletal muscle. These constructs were tested in BALB/c mice using the same paradigm. Notably, a single miR-208a binding site was sufficient to mediate potent knockdown of cardiac eGFP expression following AAV delivery (Fig. 3C).

**Fig. 3.**
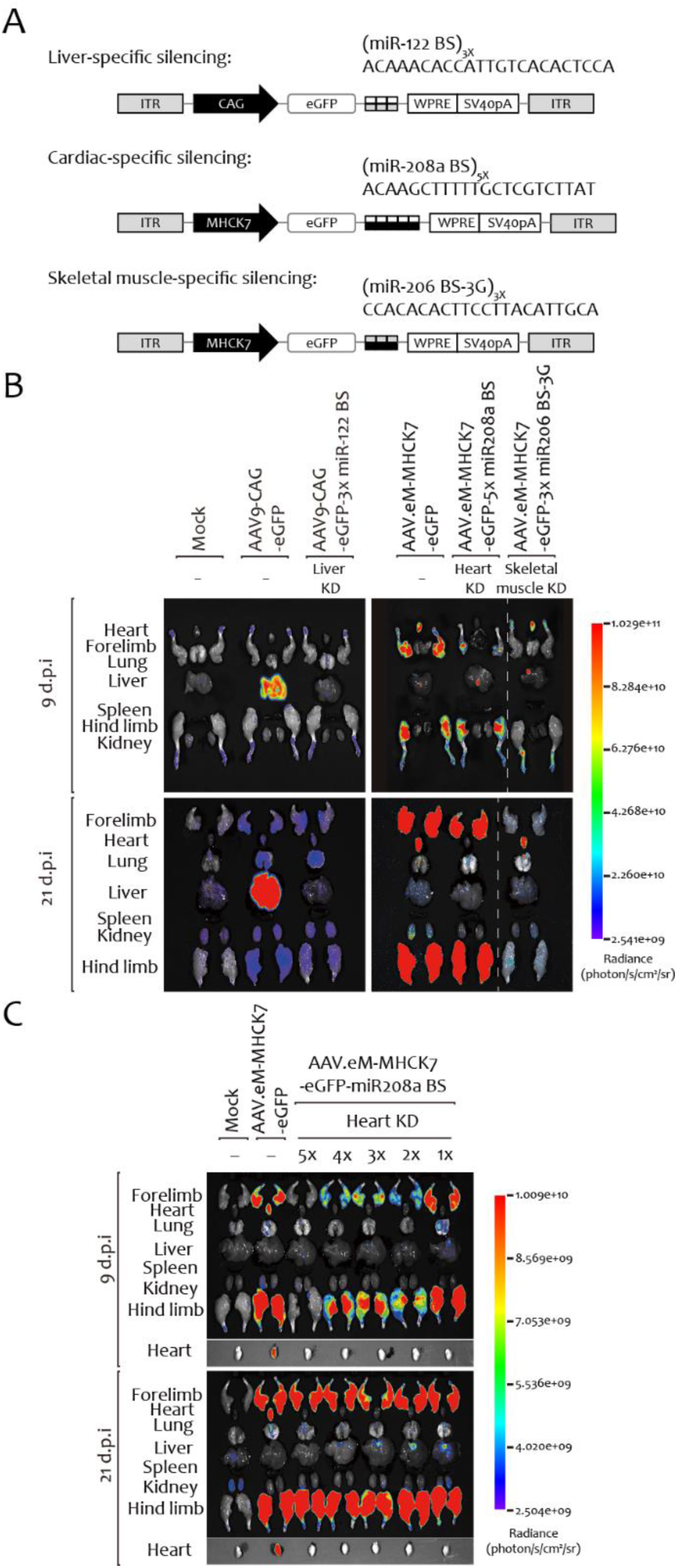
Engineering AAV vectors with tissue-selective microRNA binding sites to refine transgene expression profile. **(A)** Schematic of AAV expression constructs, detailing the identity and copy number of incorporated tissue-specific microRNA binding sites. Constructs incorporate multiple tandem binding sites (BS) for miR-122 (liver-specific repression), miR-208a (heart-specific repression), or miR-206 (skeletal muscle-specific repression) within the 3’ untranslated region of the eGFP transgene, driven by a constitutive (CAG) or muscle-specific (MHCK7) promoter. **(B)** Representative fluorescent images of tissues isolated from AAV-injected BALB/c mice at 9 or 21 days post-injection. **(C)** Fluorescent images of tissues from BALB/c mice injected with AAV.eM-MHCK7-eGFP vectors containing different copy numbers of miR-208a binding sites.

To evaluate the therapeutic potential of these findings, we incorporated a single miR-208a binding site into the AAV.eM-MHCK7-αCD19αCD3 construct to restrict αCD19αCD3 BiTE expression to skeletal muscle and limit off-target effects in cardiac tissue. The efficacy of this modified construct was compared with the parental vector in the same Raji-Luc tumor model using humanized B-NDG mice. Based on prior observations that tumor clearance was incomplete when effector cells were administered on day 16 post-engraftment, the humanization schedule was advanced to day 10 in these experiments (Fig. 4A). No drastic weight changes were observed across treatment groups, indicating that AAV administration was well tolerated (Supplementary Fig. 3A). Incorporation of the cardiac-specific microRNA binding site did not diminish the therapeutic potency of AAV.eM-MHCK7, as assessed by *in vivo* imaging (Fig. 4B). Effective tumor growth control and complete clearance were achieved irrespective of the presence of the miR-208a binding site in the 3’UTR, consistent with caliper measurements and survival data (Supplementary Fig. 3B-C). Quantitative real-time PCR analysis confirmed that incorporation of a miR-208a binding site into the transgene backbone did not alter cardiac distribution but robustly reduced transgene mRNA levels in the heart (Fig. 4C-D). Consistent with this transcriptional silencing, a concomitant and significant reduction in αCD19αCD3 BiTE protein levels was observed in the hearts of mice administered with the miR-208a binding site-modified vector (Fig. 4E). In summary, our data suggest that integrating tissue-tropic AAV capsids with microRNA-mediated regulatory circuits provides a novel strategy for precision gene therapy.

**Fig. 4.**
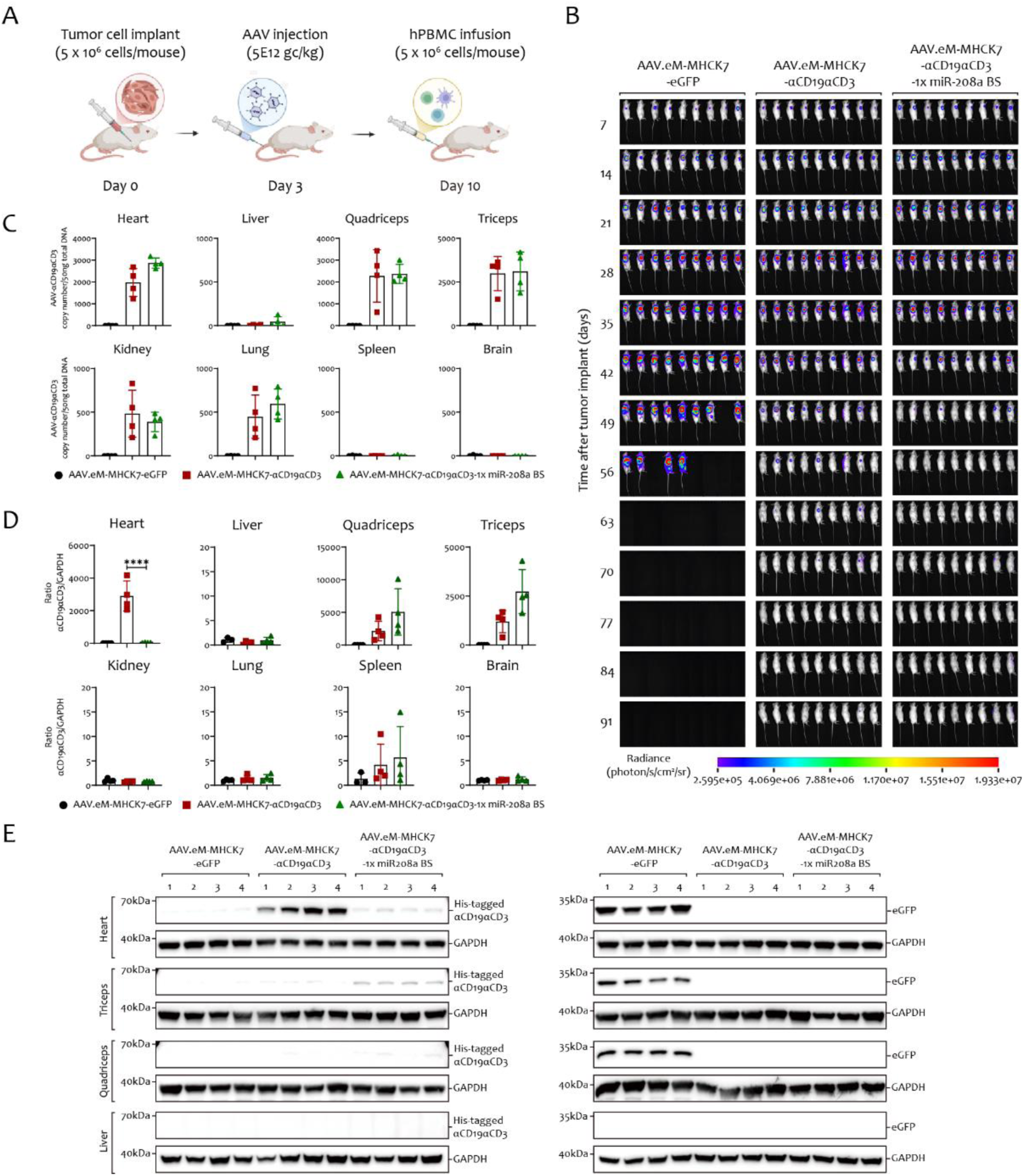
Cardiac silencing of AAV.eM-MHCK7-driven αCD19αCD3 expression using a miR-208a binding site and its impact on therapeutic efficacy and safety. **(A)** Schematic timeline of the therapeutic experiment in a humanized lymphoma model. (B) *In vivo* bioluminescent imaging of tumor burden over time in mice treated with the indicated AAV vectors. **(C)** Biodistribution of AAV vector genomes, quantified by qPCR in multiple tissues (heart, liver, quadriceps, triceps, kidney, lung, spleen, brain) collected at day 28 post tumor engraftment. Data are presented as copy numbers per 50 ng total DNA by mean ± SD (n=3-4). Statistical significance was determined by one-way ANOVA. **(D)** Quantitative RT-PCR analyses of transcript levels of the αCD19αCD3 BiTE in multiple tissues. Tissues were collected on day 28 post tumor implantation. Bar graphs show quantification of αCD19αCD3 normalized to GAPDH. Data are presented as mean ± SD (n=3-4). Statistical significance was determined by one-way ANOVA. **** *P*<0.0001. **(E)** Western blot analysis of His-tagged αCD19αCD3 and eGFP protein expression in heart, triceps, quadriceps, and liver tissues. GAPDH served as a loading control.

### Intramuscular AAV.eM injection enables 10-fold dose reduction

Systemic AAV administration can limit therapeutic efficacy and safety by pre-existing immunity, off-target transduction, and dose-limiting toxicity. Given the potential of the IM delivery to mitigate these issues, we next evaluated the therapeutic efficacy of IM-administered AAV. Immunodeficient, Raji-Luc tumor-bearing mice were first treated with AAV at a dose of 5 × 10^12^ vg/kg, following the established experimental timeline (Supplementary Fig. 4A). Both AAV9-MHCK7 and AAV.eM-MHCK7 treatments resulted in robust tumor growth restriction and clearance (Supplementary Fig4. B-D). Notably, mice treated with AAV.eM-MHCK7 exhibited significantly higher serum concentrations of the αCD19αCD3 BiTE protein compared with those receiving AAV9-MHCK7 (Supplementary Fig. 4E), indicating superior transduction efficiency of AAV.eM following IM injection. Administration of AAV.eM was also better tolerated, as reflected by a narrower range of body weight fluctuation relative to AAV9 (Supplementary Fig. 4F). No treatment-related toxicity was detected in serum cytokine and chemokine profiles measured on day 28 post AAV-injection (Supplementary Fig. 4G). Biodistribution analysis confirmed enhanced muscle tropism for AAV.eM compared with AAV9 (Supplementary Fig. 5A). Consistent with the serum drug level, AAV.eM mediated substantially higher transgene expression in muscle at both the mRNA and protein levels (Supplementary Fig. 5B-C).

Given that IM injection of either AAV9 or AAV.eM expressing the αCD19αCD3 BiTE effectively treated Raji-tumor bearing mice at a dose of 5 × 10^12^ vg/kg, and considering the substantially higher serum levels of the αCD19αCD3 BiTE achieved with the AAV.eM serotype, we hypothesized that AAV.eM would enable similarly efficacious anti-tumor treatment at a lower IM dose. To test this, the experiment was repeated at a dose of 5 × 10^11^ vg/kg (Fig. 5A). At this reduced dose, both AAV9 and AAV.eM groups exhibited stable body weight curves, indicating good tolerability (Fig. 5B). As expected, treatment with AAV.eM-MHCK7-αCD19αCD3 mediated potent tumor growth restriction at this tenfold lower dose (Fig. 5C-E). In contrast, the therapeutic efficacy of AAV9 group diminished significantly as the dose was reduced. Consistent with biodistribution data, skeletal muscle from AAV.eM-treated mice exhibited markedly higher αCD19αCD3 BiTE mRNA levels (Fig. 6A-B). Although detection of the αCD19αCD3 BiTE in tissues was challenging at this low dose due to efficient secretion into the circulation, a clear trend of higher target gene expression mediated by AAV.eM compared with AAV9 was still evident (Fig. 6C). Collectively, these results demonstrate the superior efficiency and favorable safety profile of AAV.eM for IM delivery of biologics, even at reduced vector dose.

**Fig. 5.**
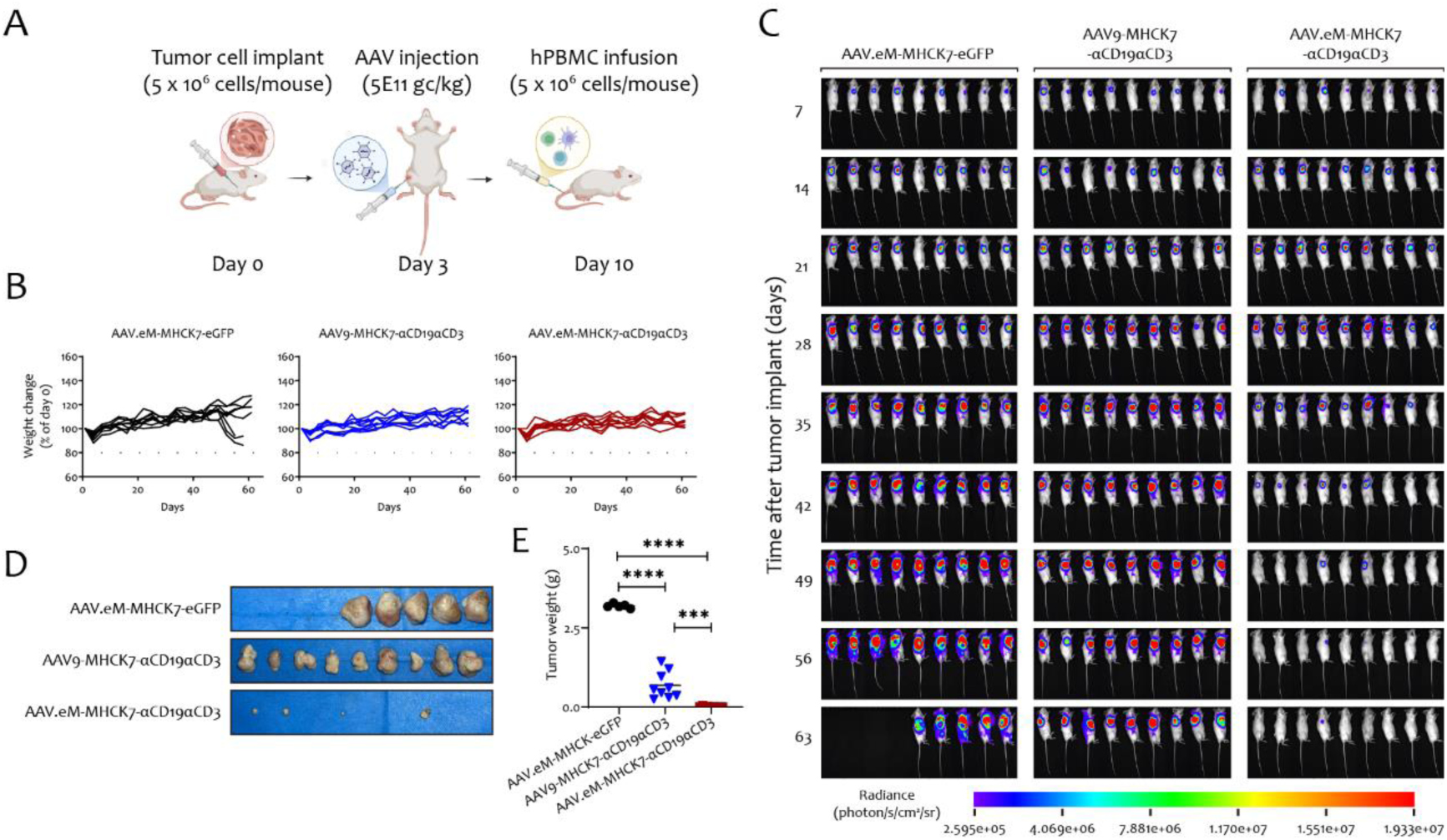
Comparative efficacy of intramuscularly administered AAV.eM and AAV9 vector for muscle-targeted αCD19αCD3 expression in a humanized lymphoma model. **(A)** Dosing regimen and treatment timeline for intramuscular injection. **(B)** Body weight change expressed as percentage of baseline (day 0) throughout the experiment. (C) *In vivo* bioluminescent imaging of Raji-Luc tumor engrafted B-NDG mice over time following treatment with the indicated AAV vector. (D) *Ex vivo* imaging of Raji tumors harvested at study termination. (E) Tumor weights correspond to specimens in panel D. Values represent the mean ± SD (n=4-9). Statistical significance was determined by one-way ANOVA. *** *P*<0.001, **** *P*<0.0001.

**Fig. 6.**
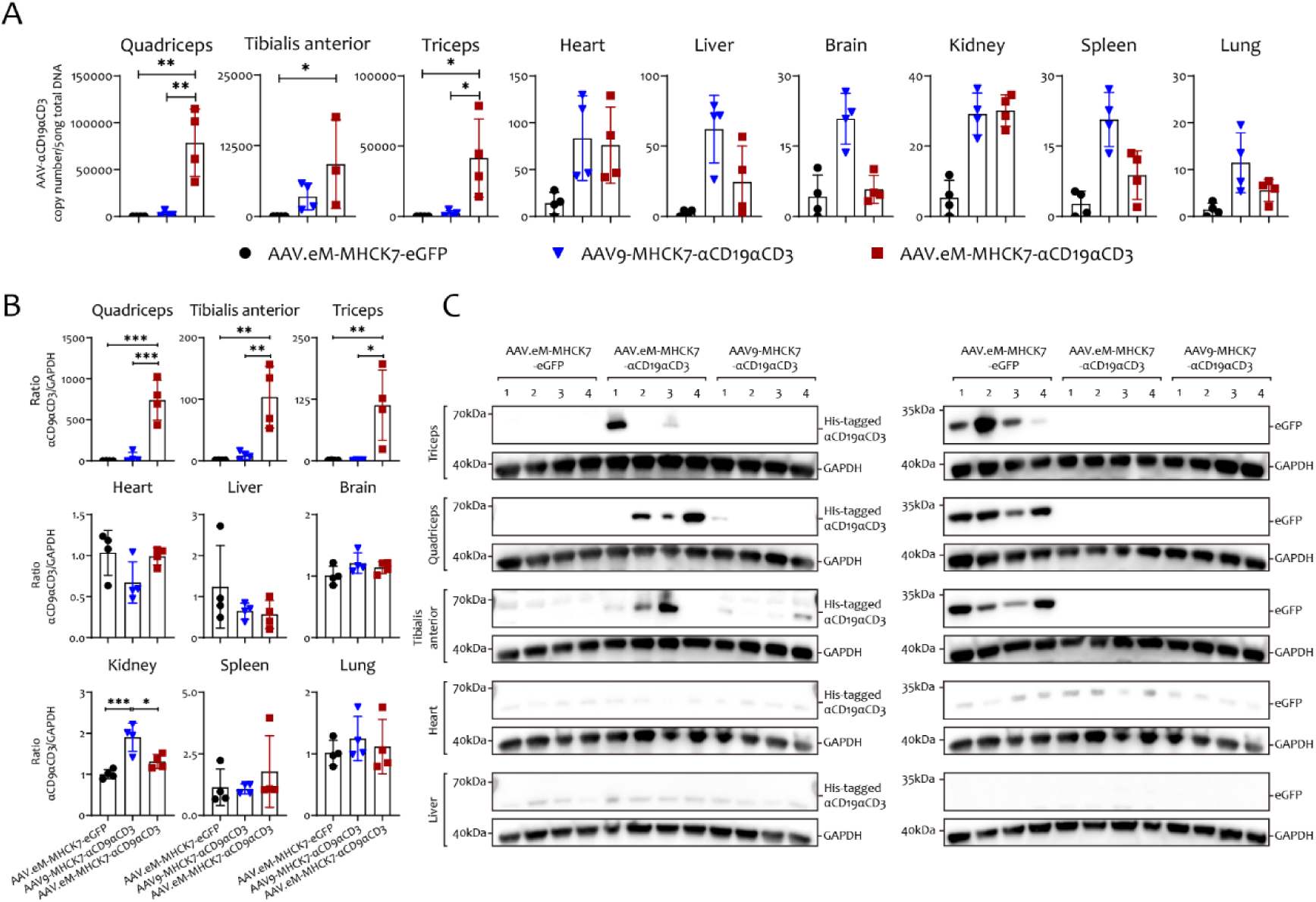
Analysis of tissue biodistribution and transgene expression following intramuscular AAV injection. **(A)** Biodistribution of AAV.eM-MHCK7 and AAV9-MHCK7 genomes in mouse tissues following intramuscular injection. AAV vector genomes were quantified by qPCR in skeletal muscles (quadriceps, triceps, tibialis anterior), heart, liver, brain, kidney, spleen, and lung collected on day 28 after tumor implantation. Bar graphs show AAV vector genome copy numbers per 50 ng total DNA at study endpoint. Data are presented as mean ± SD (n=3-4). Statistical significance was determined by one-way ANOVA. * *P*<0.05, ** *P*<0.01. **(B)** Tissue expression of αCD19αCD3 BiTE on transcript level as determined by quantitative RT-PCR. Tissues were collected on day 28 post tumor engraftment. Bar graphs represent the mean ± SD (n=3-4). Statistical significance was determined by one-way ANOVA. * *P*<0.05, ** *P<*0.01, *** *P*<0.001. **(C)** Protein expression of αCD19αCD3 BiTE intriceps, quadriceps, tibialis anterior, heart and liver tissues. GAPDH was used as a loading control.

## Discussion

The liver represents a prime target for AAV-based gene therapy owing to the pronounced hepato-tropism of most serotypes ^5^, which is facilitated by specific cell surface receptors and the organ’s unique sinusoidal architecture that promotes extensive blood contact with hepatocytes ^25^. Capitalizing on its role as a central biosynthetic platform, the transduced liver can function as an endogenous bioreactor, enabling sustained systemic production of therapeutic proteins ^26^. Effort has also been made to exploit capsid mutants with robust liver tropism ^27, 28^. However, the liver toxicity associated with AAV-mediated gene therapy has become a major concern in clinical applications, prompting extensive investigations into mechanisms of injury and risk mitigation strategies ^9^.

Skeletal muscle constitutes approximately 40% of total body mass and is predominantly composed of post-mitotic myofibers ^29^. Their long-lived nature and limited proliferative capacity enable stable, long-term transgene expression without dilution, making skeletal muscle an attractive target for sustained gene therapy ^30–32^. Kessler *et al.* demonstrated that IM administration of rAAV resulted in persistent protein expression for at least 32 weeks in myofibers ^33^. Skeletal myofibers possess the essential cellular machinery for proper antibody glycosylation, folding, and secretion ^34, 35^. Dose-dependent systematic secretion of therapeutic proteins has been observed following myotropic rAAV delivery, attributable to the high vascularity of skeletal muscle ^33^. Sustained expression of broadly neutralizing antibodies (NAbs) against simian-human immunodeficiency virus has also been demonstrated in rhesus macaques following AAV-mediated muscle transduction ^36, 37^. In the present study, we sought to alleviate AAV-associated hepatotoxicity by restricting expression of a naturally occurring serotype to muscle tissue and by comparing this approach with that of a novel muscle-selective capsid variant, AAV.eM.

A growing panel of novel AAV variants has been engineered for efficient muscle transduction. Tabebordbar *et al.* systematically identified a family of RGD-displaying capsids (MyoAAVs) that exhibit superior muscle transduction efficiency and tissue selectivity compared with AAV9 ^38^. Their lead variant retained high delivery efficacy across multiple inbred mouse strains and demonstrated translational relevance in cynomolgus macaques and human primary myotubes ^38^. Similarly, AAVMYO, an RGD motif-containing AAV9 capsid variant, mediated robust transduction cross multiple muscle types following systemic delivery while maintaining muscle-specific tropism in various mouse strains ^39^. Using a semi-rational, combinatorial bioengineering approach, the same group subsequently developed two chimeric AAV capsids, AAVMYO2 and AAVMYO3, with enhanced skeletal muscle and heart specificity, increased transduction potency, and reduced liver tropism ^40^. By integrating binding motifs specific for a skeletal muscle receptor into a liver-detargeted AAV capsid, Hong *et al.* developed LICA1, which displayed enhanced muscle tropism and superior efficacy over AAV9 in two murine models of muscular disease ^41^. Most recently, Firnberg *et al.* leveraged a combination of natural serotype screening, targeted mutagenesis, and directed evolution approach to identify several liver-detargeted, muscle-tropic AAV capsid variants ^42^.

Through capsid engineering, our group developed the novel capsid variant, AAV.eM, which exhibits enhanced myotropic properties and markedly reduced hepatic transduction^20^. The refined tissue specificity and biodistribution profile of AAV.eM have been rigorously characterized in cynomolgus monkeys. Furthermore, in a murine model of Duchenne Muscular Dystrophy (DMD), AAV.eM demonstrated superior therapeutic efficacy compared with conventional AAV9. We propose that the implementation of AAV.eM for muscle-directed gene therapy represents a promising strategy to mitigate the dose-limiting hepatotoxicity frequently associated with systemic AAV administration.

AAV9 demonstrates significantly higher transduction efficiency across many tissues and manifests a unique profile in the CNS compared with other AAV serotypes ^43–45^. In a systematic comparison of systemic delivery of AAV1–9 in mice, AAV9 exhibited rapid onset, the most widespread genome distribution, and the highest transgene expression, outperforming all other serotypes ^46^. AAV9 has thus emerged as a current “gold-standard” vector for systemic muscle-targeted gene therapy ^3^. Although challenges remain, its broad tissue tropism, superior brain blood barrier (BBB) penetration capability among WT serotypes, reduced immune clearance, and sustained gene expression establish its widespread use in clinical drug development ^47, 48^. AAV9-based platforms underpin two FDA-approved drugs and multiple active clinical trials for neuromuscular, neurodegenerative disorder, metabolic, and ophthalmic diseases ^49, 50^. Based on these considerations, AAV9 was selected as the wild-type comparator in this work.

We first evaluated our muscle-restricted αCD19αCD3 BiTE expression strategy using IV administration of AAV9-MHCK7 or AAV.eM-MHCK7 in humanized B-NDG mice bearing Raji tumors. AAV.eM out performed AAV9 in this setting, with 8 of 9 mice surviving to the endpoint and complete tumor resolution in a subset of animals. In contrast, all mice in the AAV9-MHCK7 group succumbed to the disease. The superior efficacy of AAV.eM is attributable to its enhanced muscle tropism compared with AAV9, leading to higher levels of BiTE expression and secretion into the circulation. Although AAV9 exhibits greater muscle tropism than many other wild-type serotypes, the liver remains its primary site of accumulation following systemic administration ^51, 52^. When transcription from vector genomes sequestered in the liver is not driven by a strong muscle-specific promoter, systemic transgene levels may be insufficient for therapeutic benefit. In mice treated with AAV9-CAG delivering αCD19αCD3 BiTE, we observed rapid tumor growth inhibition, consistent with the highest blood concentration among all groups. However, 6 of 9 mice experienced unexpected death after tumor regression or clearance. We detected the strongest transgene expression mediated by AAV9-CAG in the liver, in both B-NDG mice treated with αCD19αCD3 BiTE and BALB/c mice receiving dual-reporter vectors. Liver toxicity has been reported following IV administration of AAV9 in both mice and non-human primates ^53, 54^. We attempted to assess liver dysfunction by measuring the transaminase and alkaline phosphatase levels but did not detect significant differences among treatment groups, suggesting that other mechanisms may underlie the unexpected mortality observed in the AAV9-CAG group.

Given the dual tropism of AAV.eM for cardiac and skeletal muscle, we sought to develop a strategy to spatially discriminate transgene expression between these tissues. To this end, we employed a microRNA-mediated regulatory approach. Based on extensive literature review, we identified miR-208a and miR-206 as promising candidates for tissue-specific regulation ^55–59^. We engineered the transgene backbone to incorporate tandem binding sites for miR-208a or miR-206 in the 3’UTR. To prevent potential off-target repression mediated by nonspecific miR-1 binding to the miR-206 target site — potentially leading to unintended suppression of transgene expression in the heart — a single nucleotide mismatch mutation was introduced within the binding region, as reported (miR-206 BS-3G) ^21^. These designs successfully leveraged endogenous microRNA activity to achieve potent knockdown of reporter expression in the target tissue while preserving robust expression in off-target tissue. We then evaluated the therapeutic potential of these microRNA-based regulatory circuits in the same mouse model. Quantitative PCR analysis confirmed that insertion of miR-208a binding sites did not affect viral biodistribution or cardiac transduction, as evidenced by equivalent transgene DNA levels in heart tissue, but strongly reduced αCD19αCD3 BiTE transcripts in the heart. Notably, restriction of transgene expression to skeletal muscle alone did not diminish the therapeutic efficacy of AAV.eM-delivered BiTE, indicating that skeletal muscle provided sufficient production of αCD19αCD3 BiTE to achieve effective tumor control.

The strategy of harnessing tissue-specific microRNAs to suppress ectopic transgene expression in AAV-mediated gene therapy has been explored in prior studies. Qiao *et al.* attenuated transgene expression in the liver by incorporating miR-122 binding sites ^60^. Similarly, miR-122 target sequences have been used to enhance the cardiac specificity of wild-type AAV9 vectors ^61^. Other efforts have focused on mitigating transgene-directed immune response by de-targeting antigen-presenting cells (APCs) via microRNA binding sites ^62^. Our goal of differentiating transgene expression between heart and skeletal muscle using the muscle-tropic AAV capsid such as AAV.eM posed a particular challenge due to the substantial molecular similarities between these tissues ^63^. By carefully selecting tissue-specific microRNAs and optimizing tandem microRNA bindings sites in the 3’UTR, we extended the tissue-specificity of AAV.eM, achieving robust discrimination between cardiac and skeletal muscle. These findings support that incorporation of tissue-specific microRNA binding sites into vector 3’UTRs as an additional regulatory layer for precise tissue-targeted expression, further enhancing the specificity of AAV-mediated gene delivery while maintaining therapeutic efficacy.

Systemic delivery of high-dose AAV poses a risk for severe liver toxicity, as evidenced by fatal cases in real-world clinical practice, because the liver acts as a major sink for the vector uptake ^64^. IM administration may circumvent this issue, as vectors predominantly accumulate in the injected muscle with limited dissemination to other tissues, depending on serotype and dose ^33, 65–68^. Greig *et al.* demonstrated that muscle serves as the main source of systemic secretion of a transgene following IM injection, even when using ubiquitous promoters with high activity in liver and muscle ^69^. The advantages of IM over IV administration extend beyond reduced hepatic accumulation. IV-delivered AAV is highly susceptible to pre-existing neutralizing antibodies (NAbs), with titers > 1:10 often abolishing transgene expression and excluding individuals from treatment ^69, 70^. In contrast, IM administration is markedly less sensitive to pre-existing Nabs and can preserve efficient transduction ^71, 72^. In *rhesus macaques* with natural Nabs, Greig *et al.* showed that IM delivery achieved robust gene expression even at NAb titers ≤1:160 ^69^. Thus, skeletal muscle-targeted delivery significantly expands the population eligible for gene therapy compared with systemic approaches. Additionally, heterologous serotypes can be efficiently readministered to muscle without compromising expression ^69^.

Earlier studies reported rapid loss of transgene expression after IM AAV administration in large animal models, attributed to CD8+ T cells infiltration at the injection site ^73, 74^. More recent evidence indicates that this immune-mediated reduction is transient: transgene re-expression and long-term persistence (up to 5 years) have been observed, likely reflecting the stability of AAV genome in injected muscle ^75^. Similarly, long-term transgene persistence has been documented in patients receiving IM AAV1, potentially mediated by regulatory T cells (Tregs) ^76, 77^. Given the cumulative advantages of IM delivery—including reduced hepatic toxicity, lower susceptibility to pre-existing Nabs, and the possibility of redosing—the present study further investigated the therapeutic potential of AAV9-MHCK7 and AAV.eM-MHCK7 administered via IM injection.

At a dose of 5 × 10^12^ vg/kg, bilateral administration of AAV9-MHCK7 or AAV.eM-MHCK7 into the triceps, quadriceps, and tibialis anterior muscles led to substantial tumor regression or complete tumor clearance in tumor-bearing immunodeficient mice. IM delivery of both AAV9-MHCK7 and AAV.eM-MHCK7 resulted in markedly lower vector genome copies in the heart compared with IV injection at the same dose, suggesting that localized IM administration promotes highly efficient transduction within skeletal muscle while limiting systemic dissemination and off-target biodistribution.

The observation that AAV.eM-MHCK7 mediated significantly higher circulating αCD19αCD3 BiTE levels than AAV9-MHCK7 led us to hypothesize that the AAV.eM-MHCK7 dose could be reduced without compromising efficacy. Indeed, when the dose was reduced tenfold to 5 × 10^11^ vg/kg, AAV.eM-MHCK7 retained robust therapeutic efficacy, achieving efficient tumor regression or clearance, whereas efficacy of AAV9-MHCK7 was markedly diminished. These findings highlight the superior transduction efficiency of AAV.eM relative to AAV9. The ability of AAV.eM to support effective treatment at 5 × 10^11^ vg/kg has important implications for clinical translation, as it could substantially reduce manufacturing costs and lower the risk of immune-related adverse events associated with higher vector doses.

While the primary function of the AAV capsid is to determine tissue tropism and cell entry, accumulating evidence indicates that the capsid also modulates the magnitude, timing, and durability of transgene expression ^78–80^. Notably, in both our systematic and IM efficacy studies at 5 × 10^12^ vg/kg, AAV9-MHCK7 and AAV.eM-MHCK7 differed by less than 2.5-fold in viral genome delivery yet showed more than a tenfold difference in transgene levels. This discrepancy suggests that the AAV.eM capsid may confer advantages in intracellular trafficking, endosomal escape, nuclear entry, or reduced susceptibility to epigenetic silencing ^81^. These aspects merit further mechanistic investigation.

Cripe *et al.* established proof-of-principle that a single IV injection of wild-type AAV engineered to express a secreted form of blinatumomab could serve as a universal alternative to CD19-specific chimeric antigen receptor T (CAR-T) cell therapy ^82^. More recently, Song *et al.* used AAV8 combined with a liver-specific thyroxine-binding globulin (TBG) promoter to achieve hepatocyte-restricted transgene expression following systemic delivery ^83^. Employing a comparable animal model, we demonstrate here that AAV.eM-mediated, muscle-targeted transgene delivery constitutes a versatile and alternative gene therapy platform suitable for both IV and IM administration.

## Material and Methods

### Cells and cell culture

The human cell lines HEK293T and Jurkat were obtained from the American Type Culture Collection (ATCC, Manassas, VA, USA). Raji-Luc cells were purchased from Mingzhou Bio (Ningbo, China). HEK293T cells were cultured in Dulbecco’s modified Eagle medium (DMEM; Thermo Fisher, Cat#C1995500BT) supplemented with 10% fetal bovine serum (FBS; Corning, Cat#35-081-CV), 100 U/mL penicillin, and 10 μg/mL streptomycin (Biosharp, Beijing, China, Cat#BL505A). Jurkat and Raji-Luc cells were maintained in RPMI 1640 medium containing 10% FBS, 100 U/mL penicillin, and 10 mg/mL streptomycin. All cell lines were incubated at 37°C in a humidified atmosphere with 5% CO₂.

Primary human CD3+ T cells were purchased from OriCells (Shanghai, China, Cat# PB009-1F-C). Human peripheral blood mononuclear cells (hPBMCs) were obtained from OriCells (Cat# FPB004F-C) and processed according to manufacturer’s protocol for the humanization of immunodeficient mice. In brief, cryopreserved hPBMCs were thawed, transferred into a 50 mL sterile conical tube, and gently supplemented with 15-20 mL of pre-warmed RPMI 1640 medium containing a 10% FBS, 100 U/mL penicillin, and 10 mg/mL streptomycin. The cells were centrifuged at 300 × g for 10 min, and the resulting pellet was resuspended in PBS and kept on ice until further use.

### Plasmid construction and AAV viral vector production

The AAV vectors used in this study to overexpress either the αCD19αCD3 BiTE, eGFP, or the firefly luciferase 2 (Fluc2)-2A-eGFP control were custom-generated by PackGene Biotech. The constructs were engineered using overlap extension PCR, followed by Gibson assembly with homology-directed recombination. All final constructs were verified by Sanger sequencing to ensure sequence fidelity.

For viral packaging, HEK293T cells were seeded in 15 cm dishes at a density of 2 × 10⁷ cells/dish. After 24 h, the cells were co-transfected with the Rep-Cap plasmid, pHelper plasmid and the ITR-flanked transgene plasmid using polyethyleneimine (PEI). At 72 h post-transfection, cells were harvested by centrifugation (500 × g, 10 min), resuspended in PBS, and subjected to three freeze-thaw cycles to release viral particles. The crude lysate was clarified by centrifugation at 500 × g for 10 min, treated with benzonase (250 U/mL, 37°C, 30 min), and purified sequentially by iodixanol density gradient centrifugation followed by heparin sulfate affinity chromatography. Viral titers were determined by either ITR-targeted qPCR using a standard curve or TaqMan-based ddPCR. Purified AAVs were stored at −80°C with titers ranging from 1×10^12^ to 1×10^13^ vg/mL.

### Animals

Female 6-week-old non-obese diabetic CB17-Prkdcscid/l2rgtm1/Bcgen (B-NDG) mice and male 8-week-old BALB/c mice were obtained from BesTest Bio-Tech (Zhuhai, China). Animals were maintained under specific pathogen-free conditions with a 12-hour light/dark cycle and were provided with food and water *ad libitum*. All experimental procedures were conducted in accordance with protocols approved by the Institutional Animal Care and Use Committee (IACUC) of South China Normal University and the Animal Ethical and Welfare Committee of Guangzhou Forevergen Bioscience (permit no. F250423002).

### In vitro T and B cell line competition binding assays

A total of 1 × 10^6^ 293T cells were infected with AAV encoding the αCD19αCD3 BiTE at a multiplicity of infection (MOI) of 10^5^. Culture supernatants were harvested 72 hours post-infection and used for subsequent binding assays. For the binding assay, 3 × 10^5^ Jurkat (human T-cell leukemia line) or Raji (B-cell lymphoma line) cells were pre-incubated overnight with 5-20% (v/v) of the 293T supernatant. Cells were then washed with PBS, blocked with 1% human Fc blocking reagent for 10 min on ice, and stained with anti-human CD19-APC (clone HIB19, BioLegend) or anti-human CD3-FITC (clone OKT3, BioLegend) for 45 min on ice. After washing, cells were fixed in 1% paraformaldehyde (PFA) for 15 min. Flow cytometry was performed on a CytoFlex (Beckman Coulter), and data were analyzed using FlowJo.

### Killing assay

A killing assay was conducted to evaluate the ability of T cells to eliminate Raji cells in the presence or absence of αCD19αCD3 BiTE. Raji cells stably expressing luciferase (Raji-Luc) was plated into 96-well flat-bottomed plates at a density of 1 × 10^4^ cell per well. An equal volume of freshly thawed primary human CD3+ T cells (without activation) was added to Raji-Luc cells at an effector-to-target (E:T) ratio of 4:1. Each well was supplemented with 50 µL of fresh medium containing varying concentrations of clarified cell culture supernatant collected from HEK293T cells infected with an AAV vector encoding the αCD19αCD3 BiTE. The final concentration of αCD19αCD3 BiTE containing HEK293T culture supernatant was 1%, 2%, 5%, 10%, and 20%. Wells containing Raji-Luc cells and CD3+T cells without αCD19αCD3 BiTE served as negative controls, whereas wells with complete Raji-Luc lysis were used as positive controls. Following a 24-hour incubation, plates were centrifuged at 300 x g for 5 min at 4°C to separate the culture medium from cells. The luciferase signal in the clarified supernatants was measured (Beyotime Biotechnology, Shanghai, China, Cat#RG052) and normalized to the signal from the positive control. Results were expressed as the percentage of cell death.

### Enzyme-linked immunosorbent assay

IFN-γ and IL-2 produced by activated CD3+ T cells during co-culture with Raji-Luc cells and AAV-expressed αCD19αCD3 BiTE were accessed by ELISA. Briefly, Raji-Luc cells was plated into 96-well flat-bottomed plates at a density of 1 × 10^4^ cell per well. An equal volume of freshly thawed primary human CD3+ T cells (without activation) was added to Raji-Luc cells at an effector-to-target (E:T) ratio of 4:1. Each well was supplemented with 50 µL of fresh medium containing varying concentrations of clarified cell culture supernatant from HEK293T cells infected with an AAV vector encoding the αCD19αCD3 BiTE. The final concentration of αCD19-αCD3 BiTE containing HEK293T culture supernatant was 2% and 5%. Following a 24-hour incubation, plates were centrifuged at 300 x g for 5 min at 4°C to separate the culture medium from cells. The concentrations of IFN-γ and IL-2 in the supernatant were measured using ELISA kits according to the manufacturer’s instructions (BioLegend, Cat#431807 & 430107).

Serum concentrations of αCD19αCD3 BiTE in AAV-injected mice were measured using a His-tag ELISA detection kit following manufactures’ instructions (GenScript, Cat# L00436).

### Efficacy studies

Raji-Luc/GFP cells (5 × 10^6^) were implanted subcutaneously on day 0 into the right flank of 7-week-old female B-NDG mice. For IV injection, assigned cohorts of tumor-bearing mice were injected with AAVs as specified in each assay at 5 × 10^12^ gc/kg via tail vein injection on day 3. For IM injection, mice were anesthetized via intraperitoneal injection of 5% chloral hydrate (10 μL/g), achieving anesthesia within 5–10 min with a duration of approximately 1 h. Each mouse received bilateral injections (25 μL per site) into the triceps brachii, quadriceps, and tibialis anterior muscles (six sites total). The total injection volume per mouse was 150 μL. On day 10, designated cohorts were transplanted with 5 × 10^6^ hPBMCs. Mice were monitored for tumor growth and survival until a humane end point was reached. Tumor growth was evaluated by bioluminescence imaging (see below) and by caliper measurements, with tumor volume calculated using the formula: *l × w*^2^ × π/6.

### In vivo bioluminescent imaging

Longitudinal tumor monitoring was performed using bioluminescence imaging, starting 7 days post-implantation of Raji-Luc cells and continuing weekly. Prior to imaging, mice were anesthetized with isoflurane and administered Beetle D-luciferin (150 μg/g; Promega, Cat#E1605) via intraperitoneal injection. Following a 10-minute incubation period to ensure optimal substrate distribution, images were acquired using an AniView600 multi-mode in vivo animal imaging system (Biolight Biotechnology, China) with AniView Imaging software version 1.00.0044 (Biolight Biotechnology, China). Signal intensity was shown as total flux (photons/second/cm²/steradian, p/s/cm²/sr).

### Toxicity study

To evaluate systemic toxicities associated with sustained αCD19αCD3 BiTE expression, blood samples were collected in sterile 1.5 mL centrifuge tubes and allowed to clot at room temperature for 1 h on day 28 post AAV-injection. After centrifugation at 3,000 x g for 10 min at 4 °C, serum was aliquoted and stored at −80°C until analysis. Serum cytokine and chemokine profiles were determined using a cytometric bead array (CBA) system (BD Biosciences) according to manufacturer’s protocol. The following analytes were simultaneously quantified in each sample: IL-6 (Cat#558301), IL-17A (Cat#560283), IL-12/IL-23p40 (Cat#560151), IL-1α (Cat#560157), MCP-1 (Cat#558342), KC (Cat#558340), G-CSF (Cat#560152), and RANTES (Cat# 558345).

### Plasma liver biomarkers

Liver function in mice was assessed by measuring serum levels of alanine aminotransferase (ALT; Beyotime Biotechnology, Shanghai, China, Cat#P2711S), alkaline phosphatase (ALP; Beyotime Biotechnology, Shanghai, China, Cat#P0321S), aspartate aminotransferase (AST; Biosharp, Beijing, China, Cat#BL1410B) according to manufacturer’s instructions.

### Whole-tissue fluorescence imaging

Mice were anesthetized with 1% pentobarbital sodium (50 mg/kg, intraperitoneally). After anesthesia, mice were perfused with phosphate-buffered saline (PBS) via cardiac perfusion to remove blood from organs and minimize autofluorescence. The lungs, heart, spleen, kidney, liver, and limbs were collected and placed in containers on ice for imaging. Full organ imaging was performed using AniView600 multi-mode in vivo animal imaging system with AniView Imaging software version 1.00.0044 (Biolight Biotechnology). Immediately after imaging, organs were snap-frozen in liquid nitrogen and stored at −80 °C.

### Genomic DNA isolation and vector copy number analysis

Genomic DNA (gDNA) was extracted from homogenized mouse tissues using the Hipure Universal DNA Kit (Magen Biotech, Cat#D3018-03) according to the manufacturer’s protocol and quantified via Nanodrop. A standard curve was generated using a linearized transgene plasmid (αCD19αCD3 dimert sequence flanked by AAV ITRs) with a known concentration determined by droplet digital PCR (BIO-RAD). Serial dilutions (1 × 10^1^ – 1 × 10^7^ copies/μL) of the linearized plasmid were amplified in triplicate via qPCR alongside tissue gDNA. To normalize for input DNA variability, GAPDH was amplified in parallel as an internal control. Tissue gDNA/viral DNA were simultaneously assayed for both transgene and GAPDH. Primers were as follows: αCD19αCD3 dimert forward-GCTGCTGATCTACGACGC, reverse-GGGTGGATGTTCAGGGTG, probe-CATCCCCCCCAGATTCAGCGG. GAPDH forwardTGGCCTTCCGTGTTCCTAC, reverse-GAGTTGCTGTTGAAGTCGCA.

### RNA isolation and αCD19αCD3 BiTE transcripts analysis

Excised tissues were immediately preserved in RNAlater solution (Beyotime Biotechnology, Shanghai, China, Cat#R0118) and stored at −80°C until processing. Total RNA was isolated using the TransZol Up Plus RNA Kit (TransGen Biotech, Cat#ER501-01-V2) according to the manufacturer’s protocol. RNA purity and concentration were determined using a NanoDrop spectrophotometer (Thermo Fisher Scientific). For cDNA synthesis, 1 μg of total RNA was reverse transcribed using the EasyScript All-in-One First-Strand cDNA Synthesis SuperMix (TransGen Biotech, Cat#AE341-03), which includes a gDNA removal step. Quantitative PCR was performed using 2× SYBR Green qPCR Master Mix (Bimake, Cat#B21203) on a StepOne Plus Real-Time PCR System (Applied Biosystems). Primer sequences for αCD19αCD3 dimert were: forward-GGAGTGGATCGGCTACATCA, reverse-GGCAGTAGTGGTCGTCGTA. For GAPDH, the primer sequences are: forward-TGGCCTTCCGTGTTCCTAC, reverse-GAGTTGCTGTTGAAGTCGCA. Transgene expression levels were quantified using the comparative Ct (ΔΔCt) method with appropriate reference genes.

### Western blot analysis

Protein lysates were extracted from tissue samples using RIPA buffer (Beyotime Biotechnology, Shanghai, China, Cat#P0013B) supplemented with protease inhibitors (Beyotime Biotechnology, Shanghai, China, Cat#P1005B) and centrifuged at 12,000 × g for 10 min at 4°C to remove debris. Protein concentrations were determined by Bradford assay (BIO-RAD, Cat#5000205). Equal amounts of total protein (20-50 μg) were separated by SDS-PAGE on 4-12% gradient precast gels (GenScript, Cat#M00654) using MOPS running buffer (GenScript, Cat#M00680-500) at 120 V for 90 min. Proteins were transferred to PVDF membranes (Merck Milipore, Cat#ISEQ00010) using rapid transfer buffer (Beyotime Biotechnology, Shanghai, China, Cat#0572-2L) at 25 V for 30 min. Membranes were blocked with 5% non-fat milk (PHYGENE, Cat#PH1519-100g) in TBST for 1 h at room temperature and then incubated overnight at 4°C with primary antibodies against 6 × His tag (1:1000, Smart-Lifesciences, Cat#D23031507), eGFP (1:3000, Proteintech, Ca#50430-2-AP), and GAPDH (1:3000, Proteintech, Cat#10494-1-AP), diluted in antibody dilution buffer (Beyotime Biotechnology, Shanghai, China, Cat#P0023A-100mL). After three 10-min TBST washes, membranes were incubated with HRP-conjugated goat anti-rabbit secondary antibody (1:5000, Proteintech, Cat#SA00001-2) or HRP-conjugated Affipure Goat Anti-Mouse IgG (H+L) (1:5000, Proteintech, Cat#SA00001-1) for 1 h at room temperature. Protein bands were visualized using ECL substrate (Beyotime Biotechnology, Shanghai, China, Cat#P0018FS) and imaged on a ChemiDoc imaging system (BIO-RAD). GAPDH served as the loading control.

### Statistical analysis and data visualization

Statistical analyses were performed using GraphPad Prism software (Version 8; GraphPad Software). Continuous data are presented as mean ± standard deviation (SD). Differences among three or more groups were assessed by one-way analysis of variance (ANOVA). Survival distributions were compared using the Kaplan-Meier method with the log-rank test. In all analyses, a two-sided p value of less than 0.05 was considered statistically significant.

## Data availability statement

Source data for each relevant Figure is provided in a Source Data file. The data that support the findings of this study are available from the corresponding author upon reasonable request.

## Acknowledgments

We would like to acknowledge professor Guangping Gao and professor Rui Duan for their scientific input and consultancy. We are indebted to many scientists in South China Normal University and in PackGene Biotech Inc for their great assistance and collaboration.

## Author contributions

Conceptualization: H.L., Y.B., and Y.F.; Investigation: Y.F., K.T., H.C., X.C., Y.P., Y.C., and Y.A.; Methodology: Y.F., K.T., H.C., Y.P., Y.C., and Y.A.; Project administration: Y.F., and Y.B.; Resource: H.L. and Y.B.; Supervision: H.L., Y.B., and Y.F.; Validation: Y.F. and Y.B.; Visualization: Y.F.; Original draft writing: Y.F.; Review and editing: H.L., Y.B., and Y.F.

## Declaration of interest statement

H.L., Y.F., K.T., H.C., X.C., Y.P., Y.C., Y.A., and Y.B., are employees of PackGene Biotech Inc.

---

**Figure S1.**
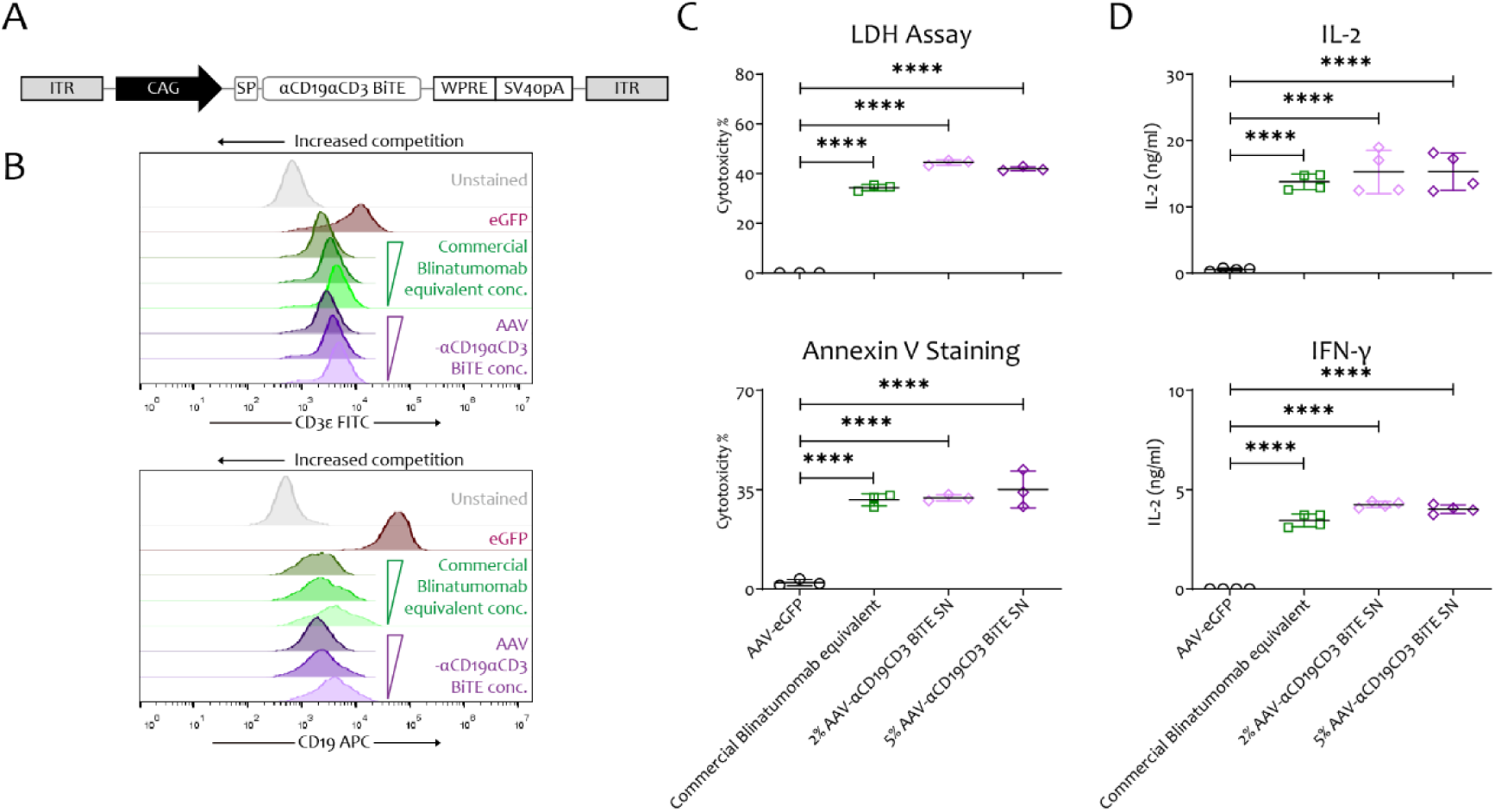
Functional characterization of AAV expressed αCD19αCD3 BiTE *in vitro*. **(A)** Schematic of the AAV BiTE expression construct. Key elements include inverted terminal repeats (ITRs), a ubiquitous promoter (CAG), a secretion signal peptide (SPI), the BiTE cassette, WPRE for enhanced expression, and an SV40 polyadenylation signal (SV40pA). **(B)** Flow cytometric analysis of Jurkat (CD3+, upper) and Raji (CD19+, lower) cells stained with fluorescent antibodies following incubation with cell culture supernatant containing AAV-derived αCD19αCD3 BiTE. **(C)** T cell-dependent cellular cytotoxicity (TDCC) of human CD3+ T cells co-cultured with Raji target cells (E:T ratio 4:1) for 48 hours in the presence of 100 ng/mL commercial Blinatumomab equivalent or titrated cell culture supernatant from AAV-BiTE-transduced cells. Cytotoxicity was quantified by measuring lactate dehydrogenase (LDH) release (top) and Annexin V staining on Raji cells (bottom). **(D)** IL-2 and IFN-γ production by CD3+ T cells upon engagement with Raji cells in the presence of indicated BiTE sources. Data are from one experiment representative of three independent experiments. Values represent the mean ± SD(n=3-4). Statistical significance was determined by one-way ANOVA. **** P<0.0001.

**Figure S2.**
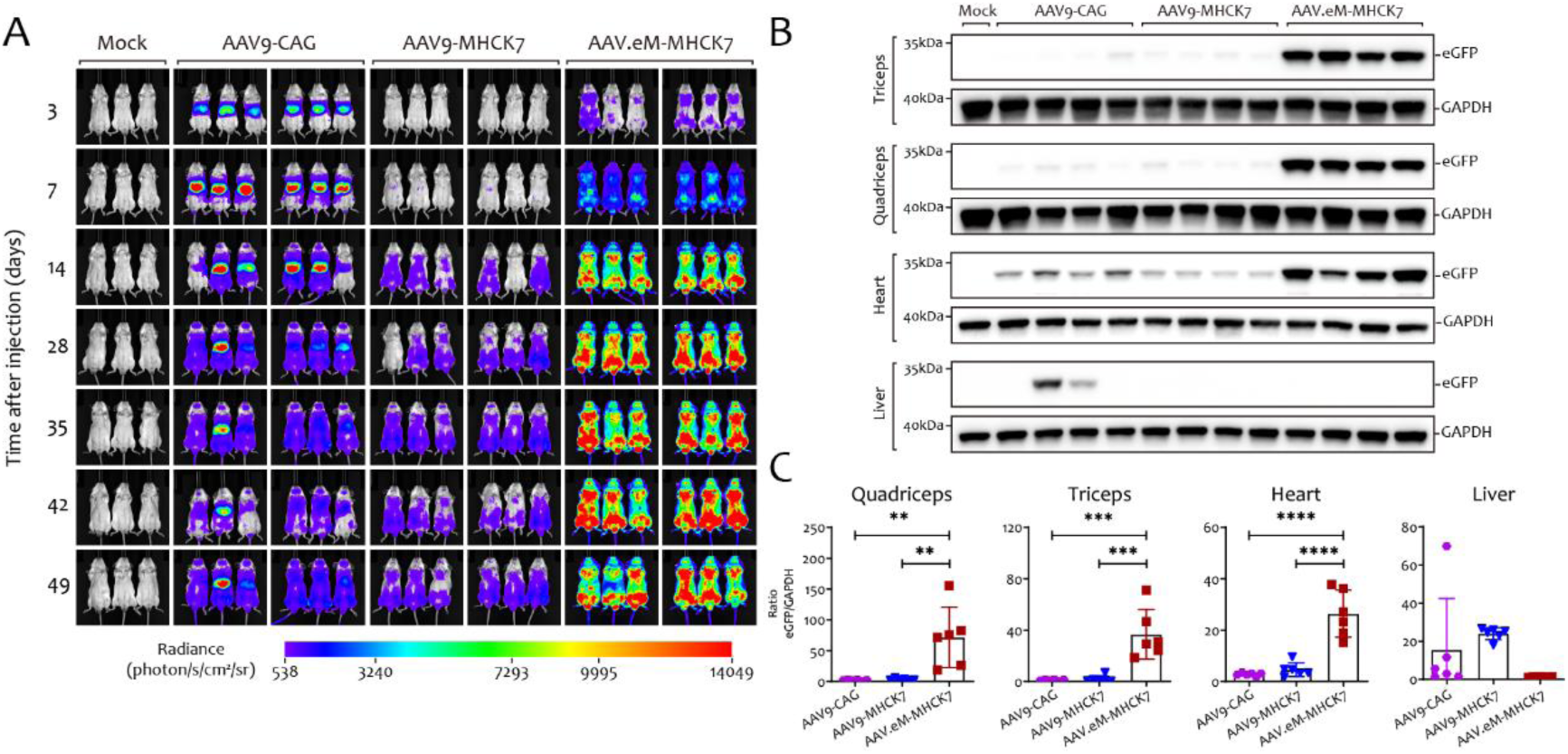
Comparative analysis of *in vivo* transduction efficiency and tissue-specific expression profiles of engineered and natural AAV vectors. **(A)** *In vivo* bioluminescent imaging of BALB/c mice following injection of AAV vectors encoding a firefly luciferase and eGFP dual reporter. Information of serotype/variant and promoter combinations were shown on the top. **(B)** Western blot analysis of tissue lysates collected from mice in (A) at day 50 post-injection. **(C)** Quantitative PCR analysis of eGFP transcript levels in tissues. Values represent the mean ± SD (n=6). Statistical significance was determined by one-way ANOVA. ** *P*<0.01, *** *P*<0.001, **** *P*<0.0001.

**Figure S3.**
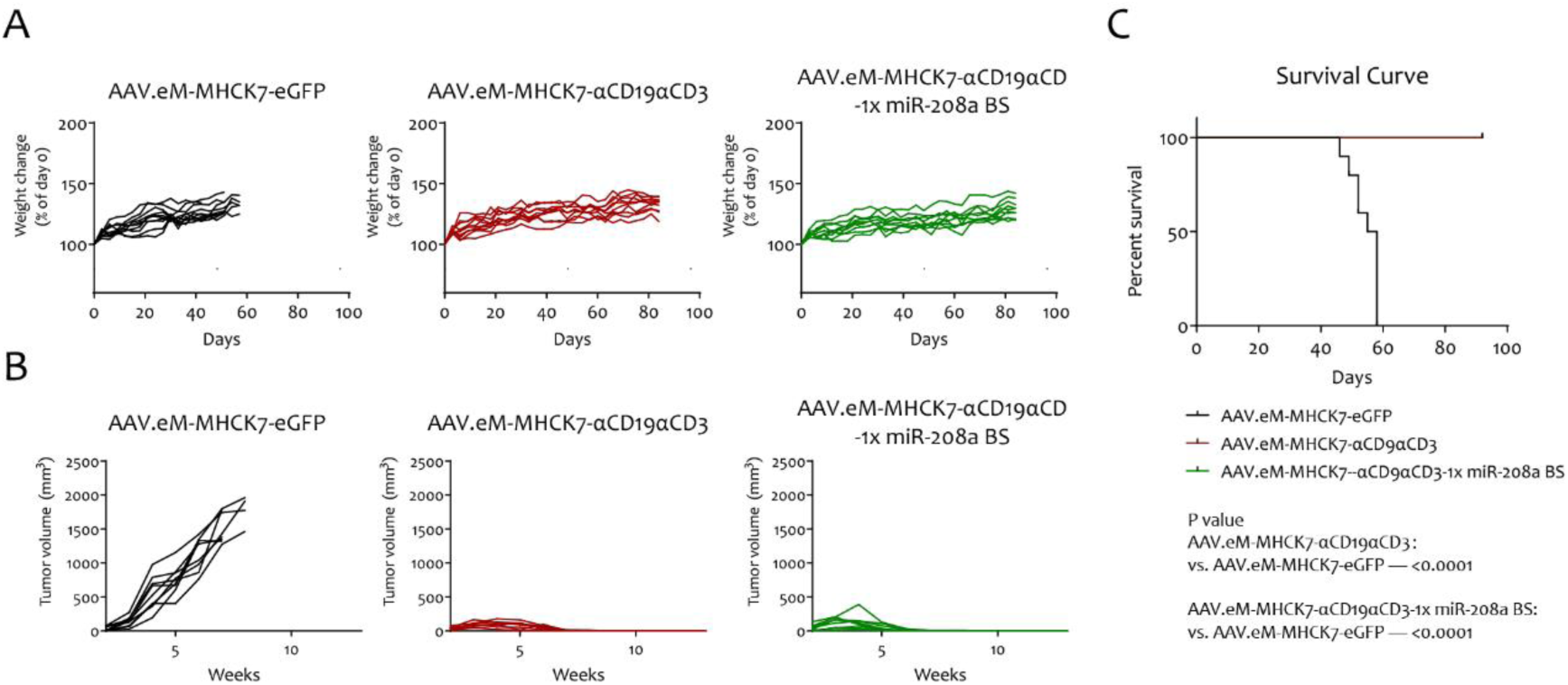
*In vivo* safety, survival, and anti-tumor efficacy of muscle-directed AAV vectors encoding αCD19αCD3 BiTE combined with miR-208a binding sites. **(A)** Individual body weights of tumor engrafted B-NDG mice, expressed as percent change from baseline. **(B)** Tumor growth kinetics measured by caliper. **(C)** Kaplan-Meier survival curve. Log-rank test was used to determine the statistical significance between groups. P value was shown in the panel. [n = 9 for AAV.eM-MHCK7-eGFP treated mice, n=9 for AAV.eM-MHCK7-αCD19αCD3 treated mice, n=9 for AAV.eM-MHCK7-αCD19αCD3-1x miR-208a BS treated mice].

**Figure S4.**
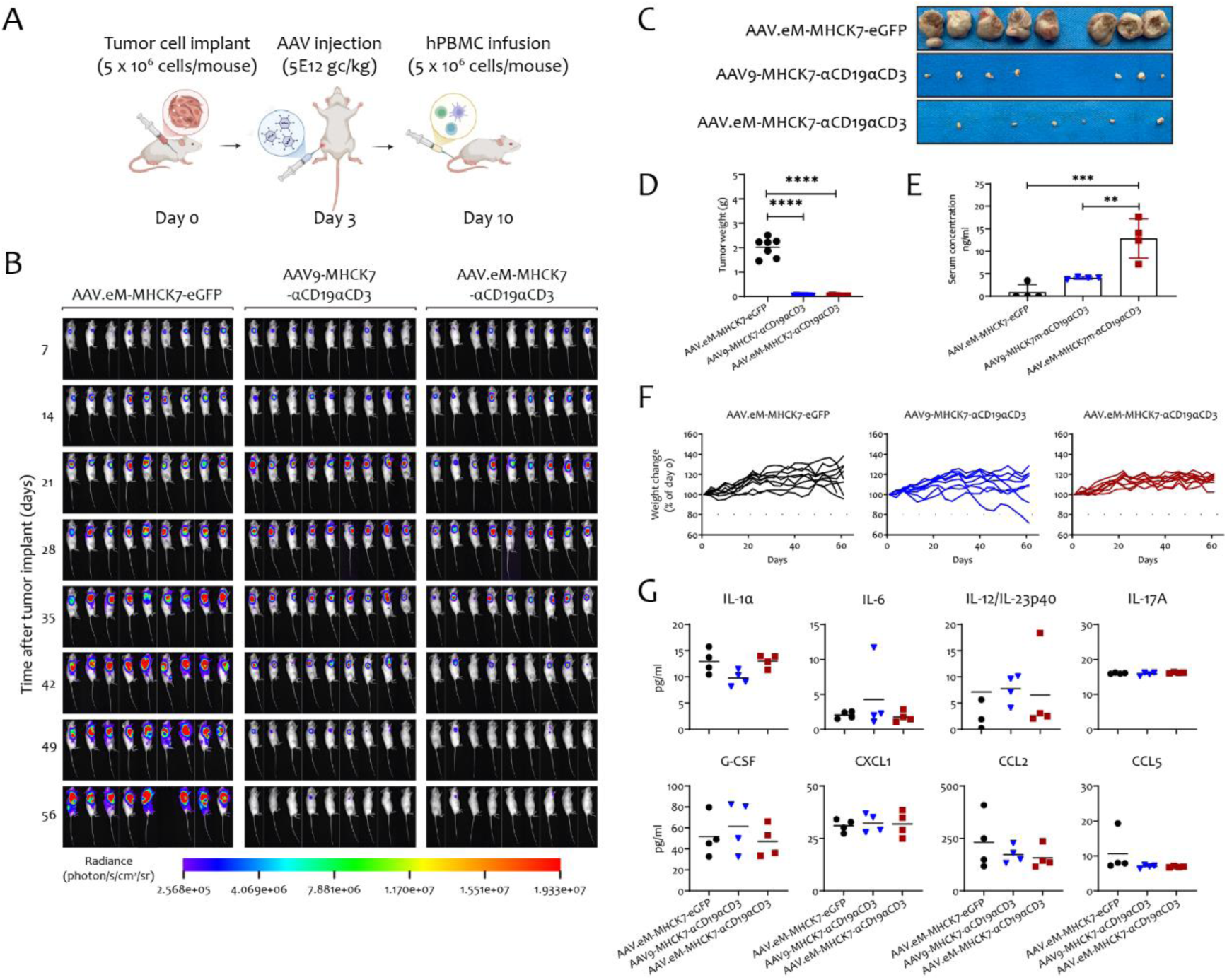
High-dose proof-of-concept for intramuscular AAV-BiTE therapy. **(A)** Experimental timeline for intramuscular injection. **(B)** *In vivo* bioluminescent imaging of individual mice bearing Raji-Luc tumors. [n = 13 for AAV.eM-MHCK7-eGFP treated mice. n= 13 for AAV9-MHCK7-αCD19αCD3 treated mice. n= 13 for AAV.eM-MHCK7-αCD19αCD3 treated mice. n=4 from each group were used for the detection of αCD19αCD3 BiTE, cytokines and chemokines in the serum.] **(C)** Mice body weight changes presented as percent change from initial weight. **(D)** Ex vivo images of Raji tumors collected at endpoint. **(E)** Tumor weights corresponding to samples in panel D. **(F)** Concentrations of AAV-expressed αCD19αCD3 BiTE in the serum at day 28 post tumor cell engraftment, quantified by anti-His tag ELISA. **(G)** CBA analyses of murine cytokine and chemokine levels in the serum collected at day 28 post tumor engraftment. Values represent the mean ± SD (n=4). Statistical significance was determined by one-way ANOVA. ** *P*<0.01, *** *P*<0.001, **** *P*<0.0001.

**Figure S5.**
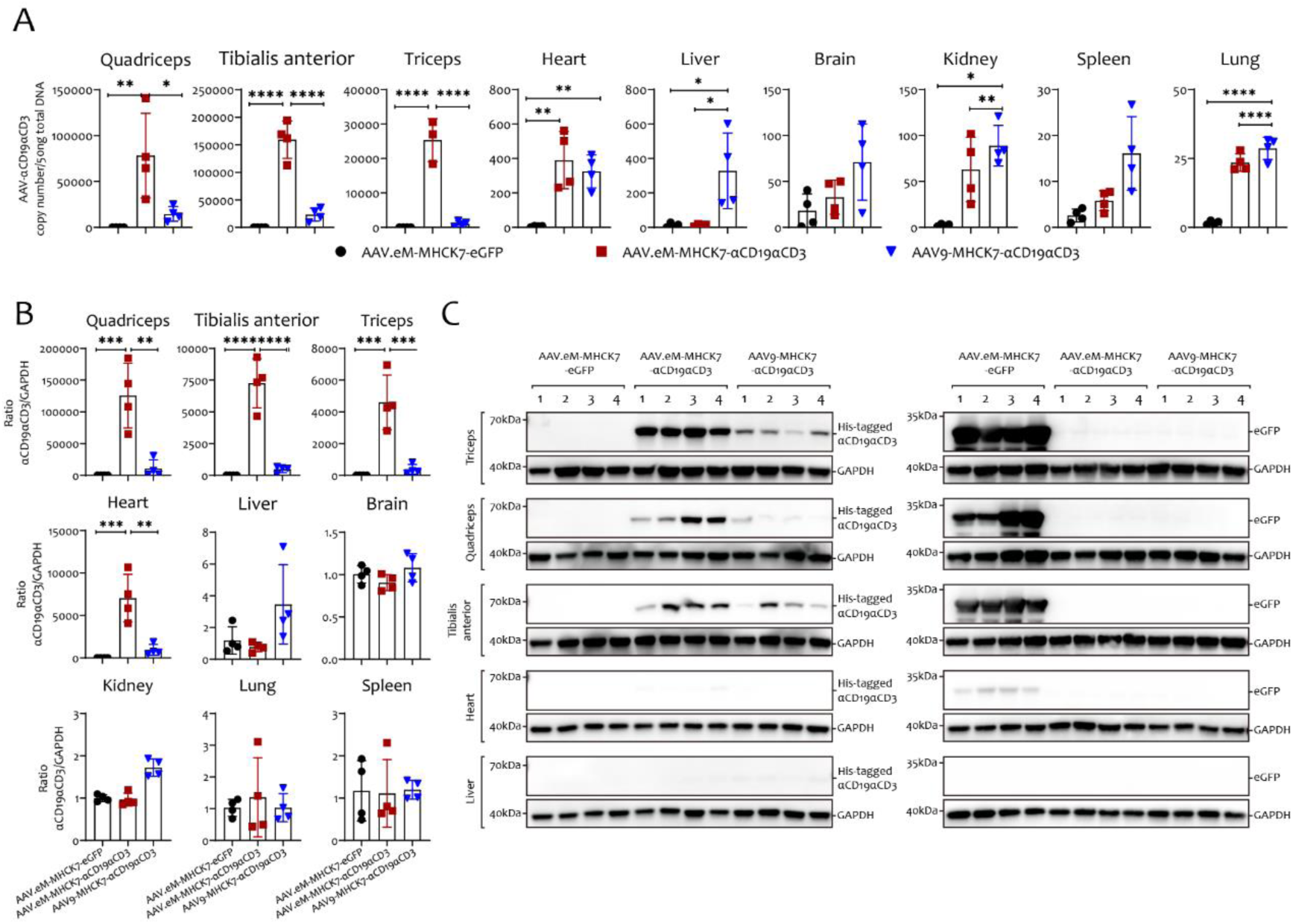
Biodistribution and tissue-specific expression of intramuscular high-dose AAV-BiTE. **(A)** Biodistribution of AAV.eM-MHCK7 and AAV9-MHCK7 genomes in mouse tissues following intramuscular injection. **(B)** Transcript levels of αCD19αCD3 BiTE in tissues, quantified by quantitative RT-PCR. **(C)** Protein expression of αCD19αCD3 BiTE in tissues, detected by western blot. Values represent the mean ± SD (n=4). Statistical significance was determined by one-way ANOVA. * *P*< 0.05, ** *P*<0.01, *** *P*<0.001, **** *P*<0.0001.

